# Microbiota of laboratory channel catfish skin mucosa and aquaria water exposed to chloramine-T trihydrate

**DOI:** 10.1101/2023.11.16.567396

**Authors:** Elliott Kittel, Tina Crosby, Brandon Kocurek, Andrea Ottesen, Charles Gieseker

## Abstract

Here, we describe the skin mucosa microbiome of channel catfish (*Ictalurus punctatus*) before and after exposure to chloramine-T. We also describe the aquaria water microbiome after the post-treatment period. These data provide a unique baseline description of skin mucosa and aquaria water microbiome from catfish reared in research aquaria.

**Disclaimer:** The views expressed in this announcement are those of the authors and do not necessarily reflect the official policy of the Department of Health and Human Services, the U.S. Food and Drug Administration, or the U.S. Government. Reference to any commercial materials, equipment, or process does not in any way constitute approval, endorsement, or recommendation by the Food and Drug Administration

## Announcement

Channel catfish (*Ictalurus punctatus*) are reared at the FDA CVM Office of Applied Science for use in Institutional Animal Care and Use Committee approved studies. A cohort of 34 catfish averaging 70 ± 27 g were used to evaluate how treatment with chloramine-T affects the microbiota of catfish skin mucosa and the microbiome of respective aquaria water. HALAMID AQUA® (chloramine-T) is an FDA approved drug for the “control of mortality in freshwater-reared warmwater finfish due to external columnaris disease associated with *Flavobacterium columnare*” (https://www.fda.gov/animal-veterinary/aquaculture/approved-aquaculture-drugs). This work generated baseline data describing how chloramine-T influences the skin mucosa microbiome of healthy catfish.

Catfish were divided equally between two 60-gallon flow-through aquaria held at 23°C. One aquarium was assigned as the chloramine-T treatment group, and the other as the control group. Catfish were procured from a commercial hatchery and reared at the research facility for 160 days. Baseline mucous samples were collected from two catfish in both aquaria by skin scrape between the dorsal fin and caudal fin, one day before chloramine-T treatment. The chloramine-T treatment was administered according to the approved label dosage (20 mg/L static bath immersion for 1 hour, 1 treatment/day for 3 consecutive days). Subsequently, skin mucus from control (n=3) and treatment (n=3) groups were sampled 1, 7, 14, 28, and 56 days following completion of treatment. Aquaria water was sampled using ultrafiltration (500 mL and 10 L samples) at a single timepoint, 56 days after chloramine-T treatment (dx.doi.org/10.17504/protocols.io.q26g78qy9lwz/v1).

Genomic DNA from catfish skin mucus was extracted using the ZymoBIOMICS Quick-DNA HMW MagBead kit (Zymo, Irvine, CA, United States). Genomic DNA from aquaria water was extracted using the Qiagen DNeasy PowerWater kit (Qiagen, Germantown, MD, United States). Libraries were prepared using the Illumina DNA Prep kit (Illumina Inc., San Diego, CA, United States). Libraries were sequenced on the Illumina NextSeq 2000 in paired end mode with 2 x 150 cycles using the Nextseq 2000 P3 300 cycle kit. Reads were screened and trimmed using Trimmomatic [1] and an average of 42 million reads per sample were annotated using an FDA in-house bacterial kmer database available on GalaxyTRAKR (http://galaxytrakr.org) [2].

Chloramine-T appeared to have little impact on bacterial taxa in skin mucosa of healthy catfish which were predominately Gram-negative bacteria. Based on linear discriminant analysis [3], only 3 taxa were significantly more abundant in the treatment group (*Pantoea sesami, Rhizobium etli*, and *Pseudomonas anguilliseptica, p<0*.*05*), while *Cellvibrionaceae* were less abundant in the control group (Figure 1a, c). A significant difference in abundance of bacterial taxa in aquaria water between the treated and control groups was observed (Figure 1d). Furthermore, many differences were observed among very low abundant taxa (< 1%) (Figure 1b).

**Figure 1.**
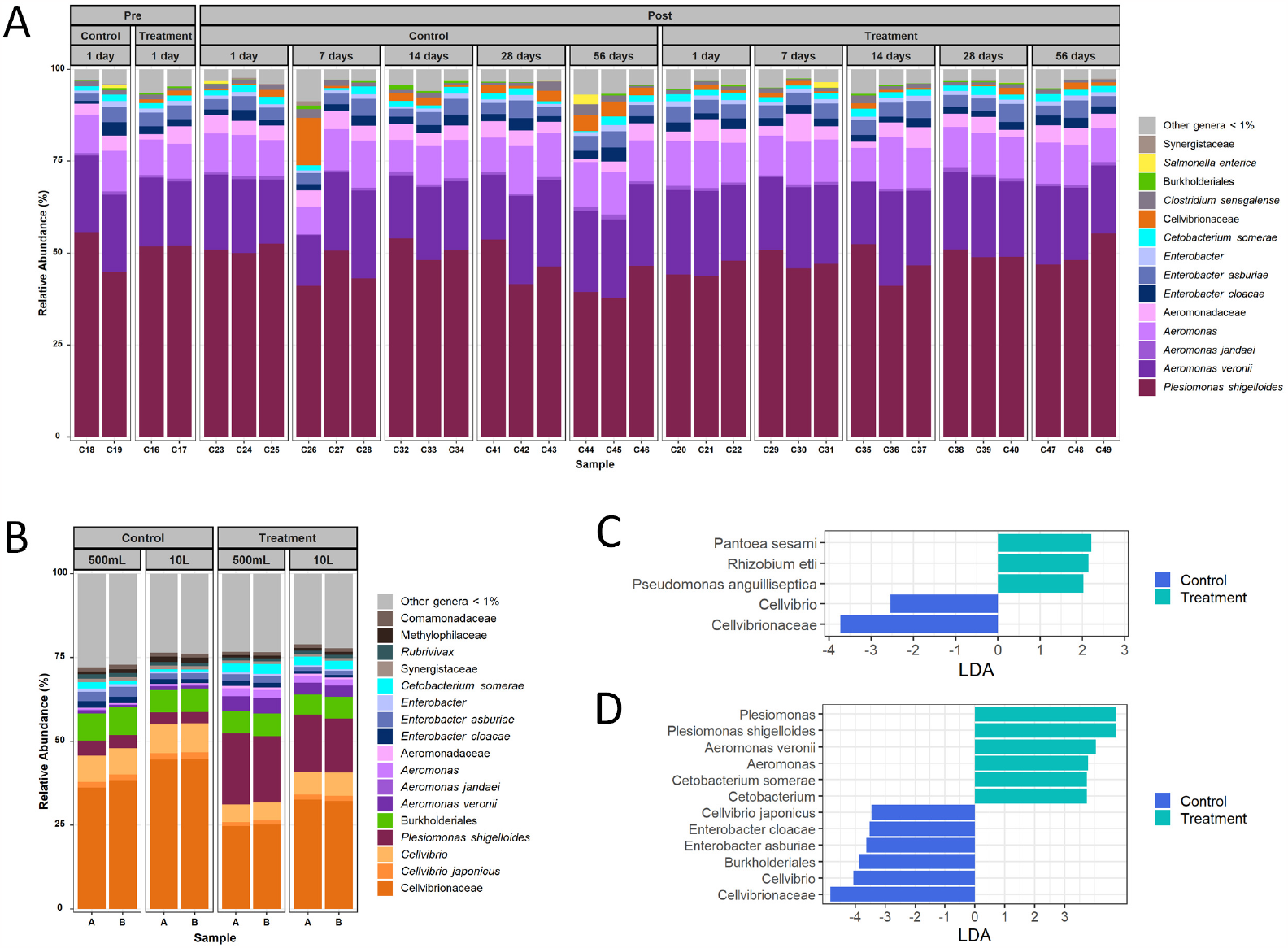
Figure 1a. Percent relative abundance of bacterial taxa associated with the catfish mucosa in control and chloramine-T treatment samples. 1b. Percent relative abundance of catfish aquaria water from 500 mL and 10 L samples in control and chloramine-T treated fish. 1c. Linear discriminate analysis (LDA) highlighting statistically significant differential abundance of specific bacterial taxa of catfish mucosa in control and chloramine-T treatment. 1d. LDA highlighting statistically significant differential abundance of bacterial taxa in control and chloramine-T treated aquaria water.

The microbiome is correlated with states of health and disease, and baseline data describing the impact of chemical treatments is valuable for supporting safe drug development [4, 5]. In this preliminary evaluation, chloramine-T did not alter the skin mucosa microbiome of healthy catfish, however additional studies are needed to better understand the effects of chloramine-T on fish microbiota.

## Data availability

The channel catfish skin mucosa data presented here have been deposited at the NCBI in BioProject PRJNA1001319 with accession numbers SAMN36808175 through SAMN36808208. The aquaria water data presented here have been deposited at the NCBI in the BioProject PRJNA1001320 with accession numbers SAMN36822970 through SAMN36822977.

## References

1. Bolger AM, Lohse M, Usadel B. 2014. Trimmomatic: a flexible trimmer for Illumina sequence data. Bioinformatics, 30(15):2114–2120.

2. Gangiredla J, Rand H, Benisatto D, Payne J, Strittmatter C, Sanders J, Wolfgang WJ, Libuit K, Herrick JB, Prarat M, Toro M, Farrell T, Strain E. 2021. GalaxyTrakr: a distributed analysis tool for public health whole genome sequence data accessible to nonbioinformaticians. BMC Genomics, 22:1–11.

3. Segata N, Izard J, Walron L, Gevers D, Miropolsky L, Garrett W, Huttenhower C. 2011. Metagenomic Biomarker Discovery and Explanation. Genome Biology, 12:R60.

4. Cho I, Blaser MJ. 2012. The human microbiome: at the interface of health and disease. Nature Reviews Genetics, 13:260.

5. Pflughoeft KJ, Versalovic J. 2012. Human Microbiome in Health and Disease. Annual Review of Pathology: Mechanisms of Disease, 7(1):99–122.

